# Biphasic bactericidal activity of nitroxoline against *Acinetobacter baumannii* isolates from urinary tract infections

**DOI:** 10.1101/2025.03.21.644696

**Authors:** Xiaofei Yi, Xin Chen, Minggui Wang, Jianfeng Zhang, Xiaogang Xu

## Abstract

*Acinetobacter baumannii* is a critical pathogen which can cause hospital-acquired infections, particularly urinary tract infections (UTIs). The antimicrobial resistance (AMR) of *A. baumannii* is rising which poses a significant challenge to clinical management. Nitroxoline, an old antibiotic for treating uncomplicated UTIs, has gained renewed interest as a potential therapeutic option. Here, we investigates the bactericidal activity of nitroxoline against 34 *A. baumannii* (17 carbapenem-resistant and 17 carbapenem-sensitive isolates) collected from UTI patients. Nitroxoline exhibited a biphasic bactericidal effect, characterized by enhanced efficacy up to an optimal bactericidal concentration (OBC), followed by declined activity at higher nitroxoline concentrations. The OBCs, minimum bactericidal concentration (MBC), minimum inhibitory concentration (MIC), and time–killing curves were evaluated to elucidate bactericidal activity. Raman deuterium stable isotope probing (Raman-DIP) was employed to validate nitroxoline’s bactericidal effect and its impact on bacterial metabolic activity and survival rates. These results demonstrate that nitroxoline exhibits excellent inhibitory and bactericidal activity against *A. baumannii*. While nitroxoline exhibits biphasic bactericidal activity with OBC_50/90_ values of 4/8 mg/L. Raman spectroscopy identified a decreased C–D ratio as nitroxoline concentration increased, indicating reduced metabolic activity. Notably, an inverse correlation (*r* = –0.7594, *p* <0.0001) was observed between bacterial survival and the ccratio at concentrations above the OBC. These findings underscore the necessity of optimizing dosing regimens to enhance nitroxoline’s therapeutic efficacy and alleviate AMR development. Raman-DIP emerges as a robust tool for assessing nitroxoline’s effects on bacterial metabolism and determining MIC, offering valuable insights for future clinical applications.

## Introduction

*Acinetobacter baumannii* has emerged as a troublesome pathogen, posing severe challenges to clinical treatment. Notably, *A. baumannii* is the fourth most common bacteria, accounting for 8.1% of infections in China (1). Urinary tract infections (UTIs) are the second most common type of infection in China, accounting for 11.65% of healthcare-associated infections(2). Among the etiologic pathogens, *A. baumannii* accounts for 1.7% of UTI cases (3). Additionally, antimicrobial resistance (AMR) in *A. baumannii* isolated from hospitals in China has notably increased. Specifically, the resistance to carbapenem which is an effective antibiotic against multi-drug resistant gram-negative pathogens, has reached 73.7% (1). Among isolates causing UTIs, the carbapenem resistance rate in *A. baumannii* is 40.3%, with extensively drug-resistant strains accounting for 16.2% of these isolates (3). Quinolones, renowned for their high concentration in urine and user-friendly application, have emerged as an essential choice for treating UTIs. However, the quinolone resistance rate in *A. baumannii* has also increased in China, with ciprofloxacin resistance reaching 43.2% in urinary isolates (3). This resistance diminishes the effectiveness of quinolones in treating UTI caused by *A. baumannii*, thereby necessitating the exploration of alternative antimicrobial agents.

Nitroxoline (5-nitro-8-hydroxyquinoline), an oral antibiotic first described in the 1950s (4), has received renewed attention for managing uncomplicated UTIs in Germany. Nitroxoline, a derivative of quinoline, is distinguished by its unique chelating mechanism of action. Nitroxoline causes microbial death by chelating and sequestering biologically important divalent ions (4). This chelation inhibits the function of bacterial RNA polymerase and biofilm formation and reduces bacterial adhesion to the bladder (5–7). However, the current clinical breakpoint of nitroxoline (susceptible, ≤ 16 mg/L), introduced by EUCAST in 2016, applies only to *Escherichia coli* and UTI (8). This is due to lower systemic concentrations than urinary concentrations observed after oral administration (standard dose 250 mg every 8 h). (9). *In vitro*, nitroxoline exhibits activity against gram-positive and gram-negative bacteria as well as fungal pathogens (10–12), demonstrating its potential for treating clinical infections.

The biphasic bactericidal activity of antibiotics is characterized by an initial increase in drug lethality at concentrations above the minimum inhibitory concentration (MIC) until the optimal bactericidal concentration (OBC) is achieved. Whereas beyond the OBC, the bactericidal activity diminishes (22,23). This phenomenon may affect clinical prescription strategies and outcomes. Biphasic responses have been widely reported for quinolones including ciprofloxacin, nalidixic acid, norfloxacin, ofloxacin, and nalidixic acid, against *E. coli* (22–25). Sparfloxacin displays a biphasic response against *E. coli*, and also *Staphylococcus aureus* and *Staphylococcus epidermidis* (23). However, the relationship between the bactericidal effect of nitroxoline against *A. baumannii* and its increasing concentration remains poorly understood. It is unclear whether nitroxoline exhibits a similar biphasic bactericidal effect.

Most previous studies on nitroxoline have determined the MIC via the broth microdilution method (BMD) (6,7,9–11), which is time-consuming and provides limited information (13). Recently, single-cell Raman spectroscopy (SCRS) has been increasingly utilized to elucidate the *in vitro* efficacy of antimicrobial agents by providing insights into bacterial metabolic activity (14–17). SCRS detects the vibration patterns of biomolecules, reflecting the biochemical characteristics or phenotypes at the single-cell level (18). Raman Deuterium Stable Isotope Probing (Raman-DIP) combines SCRS with deuterium stable isotope probing, enabling the non-destructive study of stable isotope-labeled microorganisms (19). When bacteria are cultured in heavy water (D_2_O) or deuterated substrates, active cells incorporate deuterium into their biomass via hydrogen/deuterium exchange reactions. Cells containing C−D bonds exhibit a distinguishable Raman band (2000−2300 cm^−1^), shifted from C−H vibration at ∼3000 cm^−1^. In this way, the intensity serves as a universal Raman marker for measuring the metabolic activity of single cells (14,19). Raman-DIP has often been used to determine the MIC of antibiotics (14–16). While Raman-DIP hasn’t been used for detecting bactericidal activity against *A. baumannii* by nitroxoline.

Our previous study on nitroxoline against *A. baumannii* confirmed excellent antibacterial and bactericidal activity against *A. baumannii*, with an minimum bactericidal concentration (MBC) approximately 1–2 times the MIC (21). In this study, AST and time-killing assays present the bactericidal activity of nitroxoline against *A. baumannii* was biphasic. Raman-DIP is employed for exploring *in vitro* concentration-dependent bactericidal activity of nitroxoline against *A. baumannii*. Our findings indicate that nitroxoline’s bactericidal activity operates within an OBC range, underscoring the necessity for clinicians to administer nitroxoline appropriately. Additionally, Raman-DIP has proven to be an effective tool for detection on bactericidal activity of nitroxoline against *A. baumannii*.

## Materials and methods

### Ethics

The *A. baumannii* clinical isolates used in this study were obtained from the Institute of Antibiotics strain bank at Huashan Hospital. The Ethics Committee of Huashan Hospital approved this study (approval no. KY2017-274).

### Bacterial strains

Thirty-four *A. baumannii* isolates collected from urine samples were obtained from the strain bank of the Institute of Antibiotics of Huashan Hospital in 2022. These isolates were categorized into two groups (*n* = 17/group): carbapenem-sensitive and carbapenem-resistant strains. Isolate selection was based on standard clinical microbiological procedures and diagnostic criteria for UTIs. *E. coli* ATCC 25922 was used as the quality control strain to ensure the accuracy and reliability of the experimental procedures and results.

### Antimicrobial susceptibility testing

The MICs of nitroxoline and meropenem were determined by BMD. Nitroxoline powder was dissolved in dimethyl sulfoxide (DMSO; Sangon Biotech, Shanghai, China), and dilutions were prepared in sterile water. The concentration of DMSO was adjusted to maintain consistency across all antibiotic concentrations. A bacterial suspension adjusted to 0.5 McFarland was mixed with Mueller–Hinton (MH) broth containing 0.5 to 256 mg/L antibiotics or MH broth without antibiotics. The EUCAST susceptibility breakpoint for *E. coli* (16 mg/L) was applied as a reference (8), given the lack of specific breakpoints for *A. baumannii* in the context of this study.

The MBCs and OBCs of nitroxoline were determined by plating the contents from MIC testing on Lysogeny Broth (LB) agar. A 100 μL sample was collected from the wells in the microtiter plates, where no growth was observed after a 24-h incubation at 37 °C. The liquid was washed twice with 0.85% sodium chloride solution and spread onto LB agar plates. Colonies were counted after a 24-h incubation at 37 °C. The initial number of colonies was recorded by plating the diluted inoculum solution onto LB agar. Given that the limit of detection for this technique is 10 CFU/mL, the absence of growth on the LB agar plate indicated that the concentration was < 10 CFU/mL. MBC was defined as the minimum concentration of an antimicrobial agent capable of inactivating > 99.9% of the bacteria. OBC was the concentration at which the highest bactericidal rate was achieved. Colonies that regrew were retested for MICs and compared with the original strain.

### Time–killing curve study

The time–kill kinetics of nitroxoline were determined for an *A. baumannii* strain (aba18) at concentrations from 0.5 to 256 mg/L in triplicate. Growth without nitroxoline was used as a control. Three to five single colonies were picked from LB agar plates and adjusted to a 0.5 McFarland standard. The bacterial solution was diluted in 25 mL of LB medium at a 1:100 ratio to achieve ∼10^6^ CFU/mL. After incubation at 37 °C for 0, 1, 2, 4, 8, 12, and 24 h, 1 mL of the solution was collected, and the cells were washed and serially diluted in 0.85% NaCl solution. Subsequently, 100-μL aliquots were collected and spread onto LB agar plates for viable count determination. Colonies were counted after a 24-h incubation at 37 °C.

### Raman-DIP analysis of metabolic activity in A. baumannii

Raman-DIP was used to measure the metabolic activity of five strains of *A. baumannii* after treatment with nitroxoline. The bacterial solution was diluted in MH medium using the BMD method and transferred to 96-well microplates containing 1–256 mg/L nitroxoline. After incubation at 37 °C for 2 h, D_2_O (99.9% D atoms; Sigma-Aldrich) was added to each well. The final concentration of D_2_O was 40% (v/v), allowing for rapid deuterium incorporation without significant cytotoxicity (15,19). Each plate included a positive (D_2_O) and negative (without D_2_O) control. After incubation in heavy water for 2 h, the samples were centrifuged at 10,000 rpm for 2 min. After washing with sterile water thrice, 2 μL of the suspension from each well was deposited onto an aluminum-coated slide. SCRS was acquired using a WITec confocal Raman microscope (Alpha300R, WITec, Germany) with a laser wavelength of 532 nm and a single-spectrum acquisition time of 2 s. A minimum of 20 spectra were collected from each sample. The extent of deuterium incorporation was expressed as the C−D ratio, calculated as the ratio of the C−D band intensity to the sum of the C−H and C−D band intensities (C−H band: 2800−3100 cm^−1^; C−D band: 2040−2300 cm^−1^). During Raman-DIP, the MIC of the five strains and the CFU/mL in nitroxoline-treated cultures were simultaneously measured.

### Statistical analyses

Statistical analyses and diagrams were performed using R version 4.1.1 and GraphPad Prism 6 for Windows (GraphPad Software Inc., La Jolla, CA, USA; http://www.graphpad.com). Statistical significance was set at *p* < 0.05.

## Results

### Biphasic bactericidal activity of nitroxoline against A. baumannii

The MICs, MBCs, and OBCs of nitroxoline for all isolates are presented in Table 1. Using the EUCAST susceptibility breakpoint for *E. coli* (<16 mg/L) (8), all isolates were susceptible to nitroxoline, with MIC_50/90_ values of 2/2 mg/L and a MIC range of 1–2 mg/L. Nitroxoline exhibited a low MBC_50/90_ value of 2/4 mg/L, with an MBC range of 2–8 mg/L. The MBC values were equal to or one dilution higher than the corresponding MIC values. No significant difference in nitroxoline activity was observed between the carbapenem-resistant and carbapenem-susceptible isolates. Moreover, carbapenem-resistant and carbapenem-susceptible strains had MIC_50/90_ and MBC_50/90_ values of 2/2 mg/L and 2/4 mg/L, respectively. The MIC of nitroxoline against the quality-control strain *E. coli* ATCC 25922 was 2 mg/L, within the EUCAST-defined acceptable range, whereas the MBC was 64 mg/L, five times higher than the MIC.

**Table 1.** Antimicrobial activity of nitroxoline and meropenem against *A. baumannii*.

| Strain | Carbapenem<br>resistance | Meropenem | Nitroxoline |  |  |  |  |
| --- | --- | --- | --- | --- | --- | --- | --- |
|  |  | MIC | MIC | MBC | OBC | MBC/MIC | OBC/MIC |
| aba1 | R | 32 | 2 | 2 | 2 | 1 | 1 |
| aba2 | R | 32 | 2 | 2 | 4 | 1 | 2 |
| aba3 | R | 64 | 2 | 4 | 8 | 2 | 4 |
| aba4 | R | 64 | 2 | 4 | 4 | 2 | 2 |
| aba5 | R | 32 | 1 | 2 | 8 | 2 | 8 |
| aba6 | R | 64 | 2 | 2 | 2 | 1 | 1 |
| aba7 | R | > 128 | 2 | 2 | 2 | 1 | 1 |
| aba8 | R | > 128 | 2 | 4 | 8 | 2 | 4 |
| aba9 | R | 64 | 2 | 8 | 16 | 4 | 8 |
| aba10 | R | 64 | 2 | 4 | 4 | 2 | 2 |
| aba11 | R | 16 | 2 | 4 | 4 | 2 | 2 |
| aba12 | R | 64 | 2 | 4 | 4 | 2 | 2 |
| aba13 | R | 16 | 2 | 2 | 2 | 1 | 1 |
| aba14 | R | 64 | 2 | 4 | 4 | 2 | 2 |
| aba15 | R | 32 | 2 | 2 | 4 | 1 | 2 |
| aba16 | R | 16 | 2 | 2 | 2 | 1 | 1 |
| aba17 | R | 0.5 | 2 | 2 | 2 | 1 | 1 |
| aba18 | S | 0.25 | 2 | 2 | 2 | 1 | 1 |
| aba19 | S | 0.125 | 1 | 2 | 2 | 2 | 2 |
| aba20 | S | 0.125 | 2 | 2 | 2 | 1 | 1 |
| aba21 | S | 0.125 | 1 | 2 | 2 | 2 | 2 |
| aba22 | S | 0.25 | 2 | 4 | 4 | 2 | 2 |
| aba23 | S | 0.125 | 2 | 2 | 4 | 1 | 2 |
| aba24 | S | 0.25 | 2 | 2 | 2 | 1 | 1 |
| aba25 | S | 0.25 | 2 | 2 | 4 | 1 | 2 |
| aba26 | S | 0.125 | 2 | 4 | 4 | 2 | 2 |
| aba27 | S | 0.125 | 2 | 2 | 4 | 1 | 2 |
| aba28 | S | 0.25 | 2 | 4 | 4 | 2 | 2 |
| aba29 | S | 0.125 | 2 | 2 | 2 | 1 | 1 |
| aba30 | S | 0.25 | 2 | 2 | 8 | 1 | 4 |
| aba31 | S | 0.125 | 2 | 4 | 4 | 2 | 2 |
| aba32 | S | 0.125 | 2 | 2 | 2 | 1 | 1 |
| aba33 | S | 0.015 | 1 | 2 | 2 | 2 | 2 |
| aba34 | S | 0.25 | 2 | 2 | 2 | 1 | 1 |

When testing the MBCs, nitroxoline exhibited biphasic bactericidal activity and this biphasic activity was present in both carbapenem-resistant (Figure 1a) and carbapenem-susceptible (Figure 1b) strains of *Acinetobacter baumannii*. The survival rate decreased with increasing nitroxoline concentration until the OBC was reached, which was 1- to 8-fold the MIC. Above this concentration, the survival rate increased, with an OBC_50/90_ ratio of 4/8 mg/L and a range of 2–32 mg/L (Table 1). The MICs for the regrown colonies were similar to those of the ancestral strains. Biphasic bactericidal activity was also evident in the time–killing curves for *A. baumannii* aba18 (Figure 2). The number of colonies decreased as the concentration increased during the first 4 h of testing. While the bactericidal efficiency of nitroxoline diminished with the increasing concentration above 8/16 mg/L, which was particular evident at 24 h. The OBC was observed at 8 mg/L, where nitroxoline reduced CFU by ≥ 4 log.

**Figure 1.**
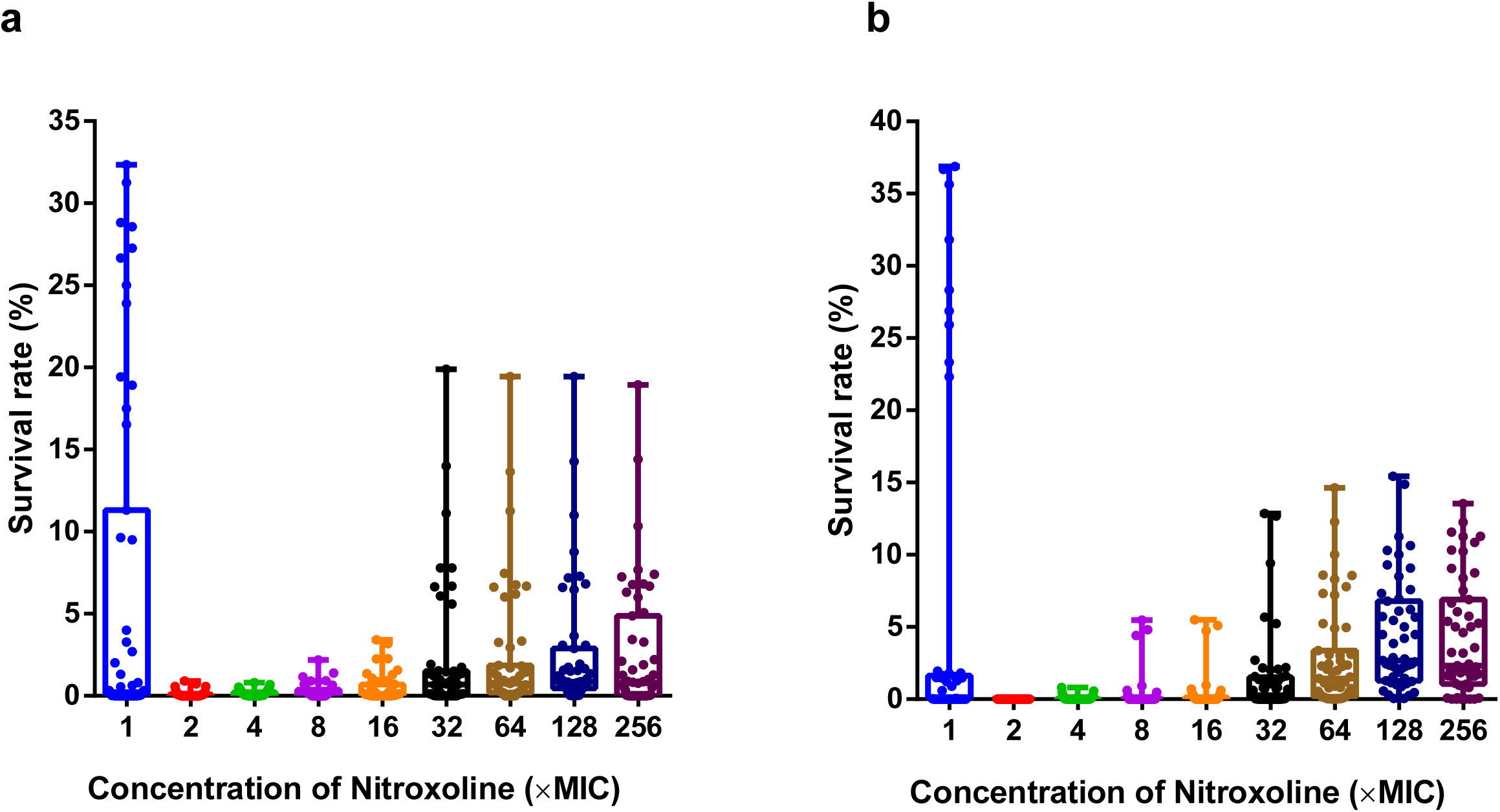
Biphasic bactericidal activity of nitroxoline. The survival of 17 strains of carbapenem-resistant *A. baumannii* (Figure 1a) *and* 17 strains of carbapenem-susceptible *A. baumannii* (Figure 1b) decreased as nitroxoline concentration increased over a low concentration range, reaching a minimum at a nitroxoline concentration of 1- to 8-fold MIC; subsequently, survival increased with nitroxoline concentration.

**Figure 2.**
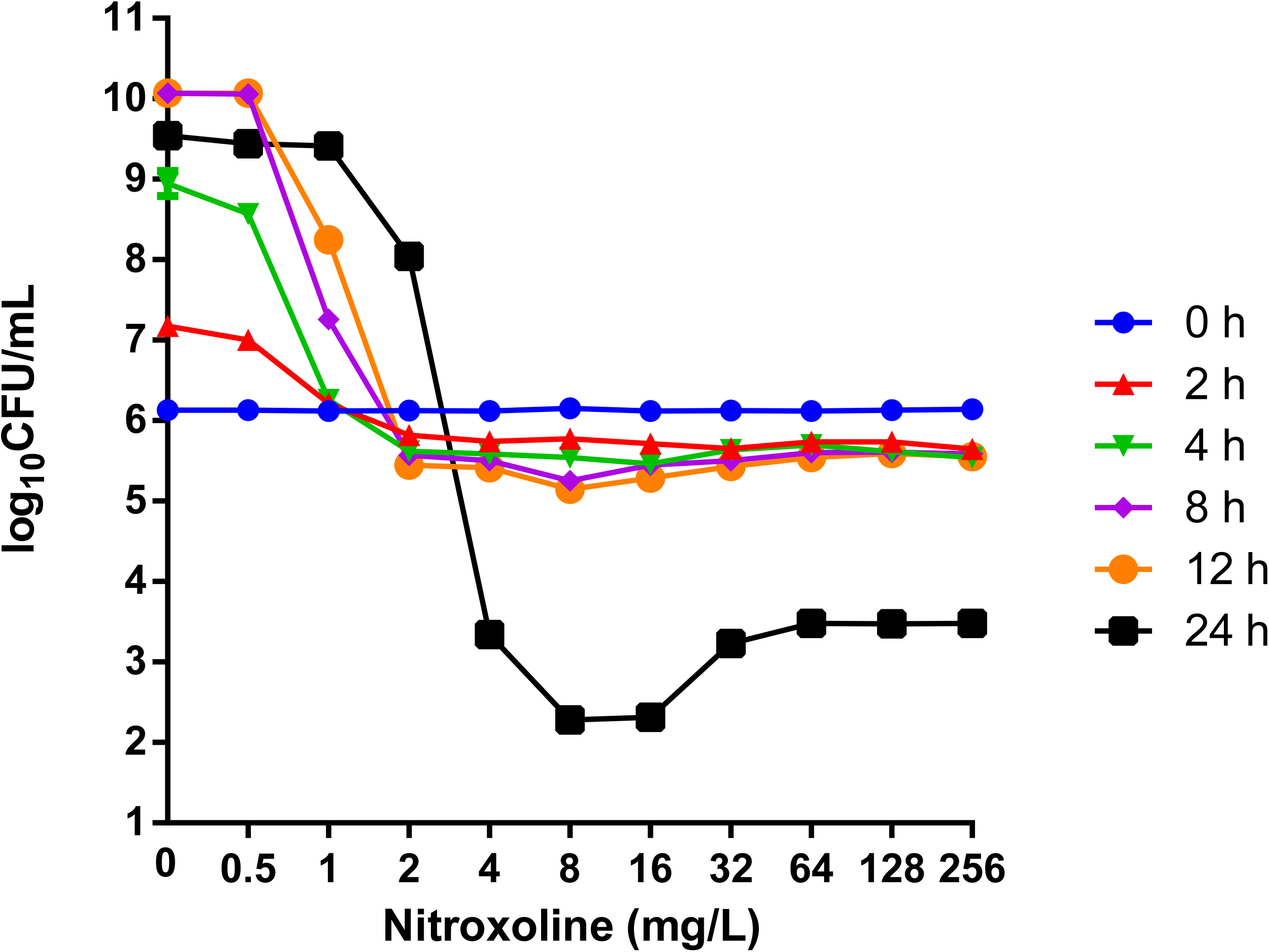
Bactericidal activity of nitroxoline against *A. baumannii* aba18 at different time points (0 h, 2 h, 4 h, 8 h, 12 h, 24 h).

### Relationship between metabolic activity and survival rate detected by Raman spectra

The deuterium incorporation in five strains of *A. baumannii* after incubation with nitroxoline and D_2_O was observed based on C−D band detection in their SCRS. The C−D bands between 2040 and 2300 cm^−1^ were identified in the SCRS in the absence of nitroxoline (Figure 3a). The C–D ratio decreased as the nitroxoline concentration increased (Figure 3b). Consistent with findings from previous studies, a 75% decrease in the C−D ratio was used to determine the MIC using Raman-DIP (23), yielding MIC values of 2 mg/L for all five strains, consistent with those obtained by BMD. At concentrations ranging from 0 to 32 mg/L, a significant difference was observed between the deuterium peaks of the bacteria and those of the negative control without the addition of D_2_O. In contrast, no significant difference was observed between the deuterium peaks of bacteria grown at concentrations of 64–256 mg/L and those of the negative control. However, some bacteria remained viable at these concentrations. Hence, despite the metabolic activity reduced significantly in the high-concentration group, *A. baumannii* strains were still able to survive. When the nitroxoline concentration exceeded the MIC, the survival rates of the five *A. baumannii* strains increased with increasing nitroxoline concentration, and their OBCs were equivalent to the MIC (2 mg/L; Figure 3c).

**Figure 3.**
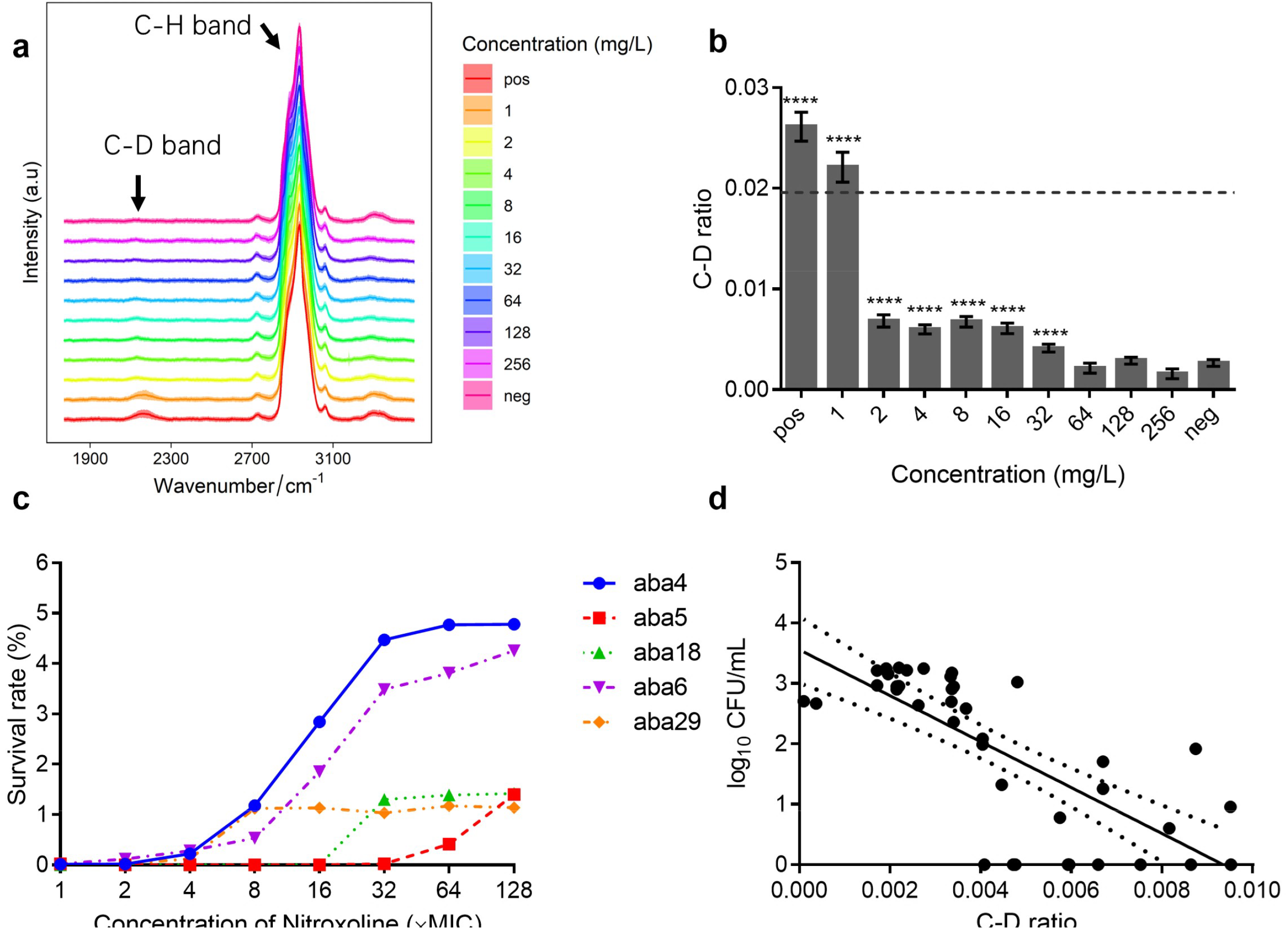
Relationship between the metabolic activity and survival rate of *A. baumannii* after nitroxoline treatment. (a) Single-cell Raman spectra (SCRS) of five *A. baumannii* isolates exposed to nitroxoline and D_2_O. (b) C−D ratio of spectra in (a); \*\**p* ≤ 0.01, \*\*\*\**p* ≤ 0.0001. The C−D ratio was defined as the percentage of the integrated spectral intensity of the C−D band (2040−2300 cm^−1^) compared to the sum of the C−D band and the predominant C−H band (2800−3100 cm^−1^); black dotted lines (75%) are the criteria to determine antibiotic susceptibility. (c) Survival rate of five *A. baumannii* isolates (MIC = 2 mg/L) after 24 h of treatment with different concentrations of nitroxoline. Survival rate was defined as the percentage of the CFU/mL after 24 h of treatment compared to the initial inoculum. (d) Inverse correlation between log_10_CFU/mL 24 h after nitroxoline addition and the C−D ratio in (b) (nitroxoline range: 2–256 mg/L); black dotted lines indicate the 95% confidence interval.

Additionally, the CFUs/mL in the cultures when the nitroxoline concentration exceeded the OBC were compared using the C−D ratio. An inverse correlation was observed between the number of surviving bacteria after treatment with nitroxoline (log_10_CFU/mL) and the C−D ratio (*r* = –0.7594, *p* < 0.0001; Figure 3d). As the intensity of the C−D ratio decreased, the log_10_CFU/mL increased, and vice versa.

## Discussion

Nitroxoline exhibites excellent *in vitro* activity against carbapenem-resistant *A. baumannii* from urinary tract specimens, with MIC_50/90_ and MBC_50/90_ values of 2/2 and 2/4 mg/L, respectively (10, 21). However, the relationship between the bactericidal effect of nitroxoline against *A. baumannii* and nitroxoline’s concentration are poorly investigated. Previous study reported that the biphasic bactericidal activity is very common in quinolones (22, 23, 25). When bacterial cells are exposed to increasing concentrations of quinolone-class antibacterials, survival drops, reaching the OBC, and then recovers to 100%. The activity of quinolone-class antibacterials decreases when bacteria reduce their growth rate, which may explain the biphasic dose response of most quinolones, producing a single maximum bactericidal concentration (28). Reactive oxygen species (ROS) dominate the lethal action of narrow-spectrum quinolone-class compounds; a decrease in ROS levels accounts for the quinolone tolerance observed at very high concentrations (24). Through detecting MIC, MBC and time-killing curves of *A. baumannii* under a wide range of concentration of nitroxolince, we confirmed the biphasic bactericidal activity of nitroxoline. Given that the regrown colonies had the same MIC as the original strains, we speculated that it may be attributed to the induction of stress responses in the bacteria at higher concentrations. This may lead to the formation of persistence or other survival mechanisms that allow a subset of the population to withstand the antibiotic challenge (24,29,30). The OBC values obtained from our study offer important insights for establishing the optimal dosing regimen of nitroxoline to enhance its bactericidal effect while reducing the risk of resistance development. The current UTI treatment guidelines recommend an oral dose of 250 mg q8h, under which the urinary concentrations of unconjugated nitroxoline (5 mg/L) (10) are comparable to those of OBC_50/90_ (4/8 mg/L) achieved in our study. Similar phenomenon was observed at 9 h in the time–killing curve for *Proteus mirabilis* ATCC 29906 and *Enterococcus faecalis* ATCC 19433 (12). However, this phenomenon was not observed in *E. coli* ATCC25922 (12). Similarly, biphasic bactericidal activity was not observed in this study against *E. coli* and *K. pneumoniae* (Figure S1). Hence, in contrast to the universal biphasic bactericidal activity of quinolines (24), the biphasic bactericidal activity of nitroxoline may be species-specific. The insights of biphasic bactericidal activity of nitroxoline warrants further investigation.

We further assessed the relationship between the metabolic activity of *A. baumannii* isolates and nitroxoline’s bactericidal acivity by Raman-DIP. Bacterial cells incorporated deuterium through metabolism, together with the C−D band can serve as a direct indicator of metabolic activity (31). In this study, the C−D ratio decreased with increasing concentrations of nitroxoline, indicating that the bacterial metabolic activity declined in a negative concentration-dependent way. This reduction in metabolic activity could be passive, meaning that the drug’s action suppresses cell metabolism (32), or it could be an active response by the bacteria, lowering their metabolic activity to survive in the presence of the drug (32,33). When the C−D band intensity decreased, the bacterial survival rate increased, and vice versa. That indicates that the metabolic activity of bacteria also has a negative correlation with the bacterial survival rate during nitroxoline treatment, possibly due to increased metabolic stress or damage. Additionally, the MIC values obtained by MHD validated the reliability of Raman spectroscopy for assessing *A. baumannii* susceptibility to nitroxoline, highlighting the potential of Raman-DIP as a effective tool for evaluating bacterial metabolic responses to nitroxoline. However, in this study, many viable bacteria with very low C–D ratios remained after exposure to high concentrations of nitroxoline. Hence, the Raman-DIP method is suitable for determining the MIC, not the MBC.

In conclusion, this study revealed nitroxoline’s biphasic bactericidal activity and successfully employed Raman-DIP to evaluate its bactericidal effect against *A. baumannii*. These findings provide a theoretical basis for optimizing nitroxoline’s use in treating *A. baumannii* infections.

## Acknowledgments

Nitroxoline powder was provided free of charge by Shanghai Yahong Medical Technology Co. LTD (Shanghai, China).

## Transparency declarations

The authors declare no conflict of interest.

## Supplementary data

Figure S1 is available as Supplementary data at AAC Online.

## References

1. Qin X, Ding L, Hao M, Li P, Hu F, Wang M. 2024. Antimicrobial resistance of clinical bacterial isolates in China: current status and trends. JAC Antimicrob Resist. 6(2):dlae052. doi: 10.1093/jacamr/dlae0522.

2. Yuan S, Shi Y, Li M, Hu X, Bai R. 2021. Trends in Incidence of Urinary Tract Infection in Mainland China from 1990 to 2019. Int J Gen Med. 14:1413–1420. doi: 10.2147/IJGM.S305358

3. Li Y, Zou M, Liu W, Yang Y, Hu F, Zhu D, Xu Y, Zhang X, Zhang F, Ji P, Xie Y, Kang M, Wang C, Fu P, Xu Y, Huang Y, Sun Z, Chen Z, Ni Y, Sun J, Chu Y, Tian S, Hu Z, Li J, Yu Y, Lin J, Shan B, Du Y, Guo S, Wei L, Zou F, Zhang H, Wang C, Hu Y, Ai X, Zhuo C, Su D, Guo D, Zhao J, Yu H, Huang X, Jin Y, Shao C, Xu X, Yan C, Wang S, Chu Y, Zhang L, Ma J, Zhou S, Zhou Y, Zhu L, Meng J, Dong F, Lü Z, Hu F, Shen H, Zhou W, Jia W, Li G, Wu J, Lu Y, Li J, Duan J, Kang J, Ma X, Zheng Y, Guo R, Zhu Y, Chen Y, Meng Q, Wang S, Hu X, Shen J, Wang R, Fang H, Yu B, Zhao Y, Gong P, Weng K, Zhang Y, Liu J, Liao L, Gu H, Jiang L, He W, Xue S, Feng J, Yue C. 2024. Changing distribution and resistance profiles of common pathogens isolated from urine in the CHINET Antimicrobial Resistance Surveillance Program, 2015–2021. Chin J Infect Chemother 24:287–299. doi:10.16718/j.1009-7708.2024.03.006

4. Petrow V, Sturgeon B. 1954. Some quinoline-5: 8-quinones. J Chem Soc 0:570–574. doi:10.1039/jr9540000570

5. Pelletier C, Prognon P, Bourlioux P. 1995. Roles of divalent cations and pH in mechanism of action of nitroxoline against Escherichia coli strains. Antimicrob Agents Chemother 39:707–713. doi:10.1128/AAC.39.3.707

6. Repac Antić D, Kovač B, Kolenc M, Karačonji IB, Gobin I, Petković Didović M. 2024. Combinatory effect of nitroxoline and gentamicin in the control of uropathogenic enterococci infections. Antibiotics (Basel) 13:829. doi:10.3390/antibiotics13090829

7. Puértolas-Balint F, Warsi O, Linkevicius M, Tang P-C, Andersson DI. 2020. Mutations that increase expression of the EmrAB-TolC efflux pump confer increased resistance to nitroxoline in Escherichia coli. J Antimicrob Chemother 75:300–308. doi:10.1093/jac/dkz434

8. EUCAST. Nitroxoline: rationale for the clinical breakpoints, version 1.0, 2016. https://www.eucast.org/fileadmin/src/media/PDFs/EUCAST_files/Rationale_documents/Nitroxoline_Rationale_Document_1.0_20161111.pdf

9. Wijma RA, Huttner A, Koch BCP, Mouton JW, Muller AE. 2018. Review of the pharmacokinetic properties of nitrofurantoin and nitroxoline. J Antimicrob Chemother 73:2916–2926. doi:10.1093/jac/dky255

10. Fuchs F, Becerra-Aparicio F, Xanthopoulou K, Seifert H, Higgins PG. 2022. In vitro activity of nitroxoline against carbapenem-resistant Acinetobacter baumannii isolated from the urinary tract. J Antimicrob Chemother 77:1912–1915. doi:10.1093/jac/dkac123

11. Kresken M, Körber-Irrgang B. 2014. In vitro activity of nitroxoline against Escherichia coli urine isolates from outpatient departments in Germany. Antimicrob Agents Chemother 58:7019–7020. doi:10.1128/AAC.03946-14

12. Sobke A, Makarewicz O, Baier M, Bär C, Pfister W, Gatermann SG, Pletz MW, Forstner C. 2018. Empirical treatment of lower urinary tract infections in the face of spreading multidrug resistance: in vitro study on the effectiveness of nitroxoline. Int J Antimicrob Agents 51:213–220. doi:10.1016/j.ijantimicag.2017.10.010

13. Kumar D, Bhattacharyya S, Gupta P, Banerjee G, Singh M. 2015. Comparative analysis of disc diffusion and E-test with broth micro-dilution for susceptibility testing of clinical candida isolates against amphotericin B, fluconazole, voriconazole and caspofungin. J Clin Diagn Res 9:DC01-4. doi:10.7860/JCDR/2015/14119.6735

14. Yi X, Song Y, Xu X, Peng D, Wang J, Qie X, Lin K, Yu M, Ge M, Wang Y, Zhang D, Yang Q, Wang M, Huang WE. 2021. Development of a Fast Raman-Assisted Antibiotic Susceptibility Test (FRAST) for the antibiotic resistance analysis of clinical urine and blood samples. Anal Chem 93:5098–5106. doi:10.1021/acs.analchem.0c04709

15. Tao Y, Wang Y, Huang S, et al. 2017. Metabolic-activity-based assessment of antimicrobial effects by D2O-labeled single-cell Raman microspectroscopy. Anal Chem 89:4108–4115. doi:10.1021/acs.analchem.6b05051

16. Zhang M, Hong W, Abutaleb NS, Li J, Dong P-T, Zong C, Wang P, Seleem MN, Cheng J-X. 2020. Rapid determination of antimicrobial susceptibility by stimulated Raman scattering imaging of D_2_O metabolic incorporation in a single bacterium. Adv Sci (Weinh) 7:2001452. doi:10.1002/advs.202001452

17. Yang K, Li H-Z, Zhu X, Su J-Q, Ren B, Zhu Y-G, Cui L. 2019. Rapid antibiotic susceptibility testing of pathogenic bacteria using heavy-water-labeled single-cell Raman spectroscopy in clinical samples. Anal Chem 91:6296–6303. doi:10.1021/acs.analchem.9b01064

18. Wang D, He P, Wang Z, Li G, Majed N, Gu AZ. 2020. Advances in single cell Raman spectroscopy technologies for biological and environmental applications. Curr Opin Biotechnol 64:218–229. doi:10.1016/j.copbio.2020.06.011

19. Berry D, Mader E, Lee TK, Woebken D, Wang Y, Zhu D, Palatinszky M, Schintlmeister A, Schmid MC, Hanson BT, Shterzer N, Mizrahi I, Rauch I, Decker T, Bocklitz T, Popp J, Gibson CM, Fowler PW, Huang WE, Wagner M. 2015. Tracking heavy water (D2O) incorporation for identifying and sorting active microbial cells. Proc Natl Acad Sci USA 112: E194–E203. doi:10.1073/pnas.1420406112

20. Brauner A, Fridman O, Gefen O, Balaban NQ. 2016. Distinguishing between resistance, tolerance and persistence to antibiotic treatment. Nat Rev Microbiol 14:320–330. doi:10.1038/nrmicro.2016.34

21. Yi X, Chen X, Lu Y, Zhang J, Chen J, Wang M, Xu X. 2025. In Vitro antimicrobial activity of nitroxoline against uropathogens isolated from China. JAC Antimicrob Resist 1:dlaf012. doi:10.1093/jacamr/dlaf012.

22. Lewin CS, Morrissey I, Smith JT. 1991. The mode of action of quinolones: the paradox in activity of low and high concentrations and activity in the anaerobic environment. Eur J Clin Microbiol Infect Dis 10:240–248. doi:10.1007/BF01966996

23. Lewin CS, Morrissey I, Smith JT. 1992. The bactericidal activity of sparfloxacin. J Antimicrob Chemother 30:625–632. doi:10.1093/jac/30.5.625

24. Luan G, Hong Y, Drlica K, Zhao X. 2018. Suppression of reactive oxygen species accumulation accounts for paradoxical bacterial survival at high quinolone concentration. Antimicrob Agents Chemother 62:e01622–17. doi:10.1128/AAC.01622-17

25. Smirnova GV, Oktyabrsky ON. 2018. Relationship between Escherichia coli growth rate and bacterial susceptibility to ciprofloxacin. FEMS Microbiol Lett 365:fnx254. doi:10.1093/femsle/fnx254

26. Kranz J, Schmidt S, Lebert C, Schneidewind L, Vahlensieck W, Sester U, Fünfstück R, Helbig S, Hofmann W, Hummers E, Kunze M, Kniehl E, Naber K, Mandraka F, Mündner-Hensen B, Schmiemann G, Wagenlehner FME. 2017. [Epidemiology, diagnostics, therapy, prevention and management of uncomplicated bacterial outpatient acquired urinary tract infections in adult patients: update 2017 of the interdisciplinary AWMF S3 guideline]. Urologe A 56:746–758. doi:10.1007/s00120-017-0389-1

27. Bulens SN, Yi SH, Walters MS, Jacob JT, Bower C, Reno J, Wilson L, Vaeth E, Bamberg W, Janelle SJ, Lynfield R, Snippers Vagnone P, Shaw K, Kainer M, Muleta D, Mounsey J, Dumyati G, Concannon C, Beldavs Z, Cassidy PM, Phipps EC, Kenslow N, Hancock EB, Kallen AJ. 2018. Carbapenem-nonsusceptible Acinetobacter baumannii, 8 US Metropolitan Areas, 2012–2015. Emerg Infect Dis 24:727–734. doi:10.3201/eid2404.171461

28. Baquero F, Levin BR. 2021. Proximate and ultimate causes of the bactericidal action of antibiotics. Nat Rev Microbiol 19:123–132. doi:10.1038/s41579-020-00443-1

29. Bakkeren E, Diard M, Hardt W-D. 2020. Evolutionary causes and consequences of bacterial antibiotic persistence. Nat Rev Microbiol 18:479–490. doi:10.1038/s41579-020-0378-z

30. Niu H, Gu J, Zhang Y. 2024. Bacterial persisters: molecular mechanisms and therapeutic development. Signal Transduct Target Ther 9:174. doi:10.1038/s41392-024-01866-5

31. Ueno H, Kato Y, Tabata KV, Noji H. 2019. Revealing the metabolic activity of persisters in mycobacteria by single-cell D2O Raman imaging spectroscopy. Anal Chem 9:15171–1518. doi:10.1021/acs.analchem.9b03960

32. Lobritz MA, Belenky P, Porter CBM, Gutierrez A, Yang JH, Schwarz EG, Dwyer DJ, Khalil AS, Collins JJ. 2015. Antibiotic efficacy is linked to bacterial cellular respiration. Proc Natl Acad Sci USA 112:8173–8180. doi:10.1073/pnas.1509743112

33. Meylan S, Porter CBM, Yang JH, Belenky P, Gutierrez A, Lobritz MA, Park J, Kim SH, Moskowitz SM, Collins JJ. 2017. Carbon sources tune antibiotic susceptibility in Pseudomonas aeruginosa via tricarboxylic acid cycle control. Cell Chem Biol 24:195–206. doi:10.1016/j.chembiol.2016.12.015

